# Free-Form Microfluidic Microneedle Array Patches

**DOI:** 10.1101/2025.02.03.635973

**Authors:** Ian A. Coates, Madison M. Driskill, Netra U. Rajesh, Gabriel Lipkowitz, Dan Ilyin, Yue Xu, Maria T. Dulay, Gunilla B. Jacobson, Jillian L. Perry, Shaomin Tian, Joseph M. DeSimone

**Affiliations:** Department of Chemical Engineering, Stanford University, Stanford, CA 94305, USA; Department of Bioengineering, Stanford University, Stanford, CA 94305, USA; Department of Mechanical Engineering, Stanford University, Stanford, CA 94305, USA; Department of Radiology, Stanford University, Stanford, CA 94305, USA; Center for Nanotechnology in Drug Delivery and Division of Pharmacoengineering and Molecular Pharmaceutics, Eshelman School of Pharmacy, University of North Carolina at Chapel Hill, NC 27599, USA; Department of Microbiology and Immunology, University of North Carolina at Chapel Hill, Chapel Hill, NC 27599, USA

## Abstract

Personalized biomedical devices, such as microneedle array patches (MAPs), offer a promising transdermal drug delivery technology, providing a safe, painless, and self-administered alternative to traditional hypodermic injections. Despite their potential for precise therapeutic release, MAP adoption has been limited by challenges in payload capacity, treatment versatility, and manufacturing scalability. To address these issues, we integrated microfluidic channel designs with MAP technology, enhancing its functionality for delivering a range of payloads, from liquid therapeutics to solid-state cargos, at tunable volumes. Using injection continuous liquid interface production (iCLIP), a novel additive manufacturing approach, we fabricated high-resolution microfluidic MAPs with complex designs. Inspired by the stingers and fangs of various venomous animals, we developed a biomimetic microneedle design that prevents clogging, enhances mechanical strength, and eliminates needle leakage, thereby improving therapeutic delivery efficiency. Our technology reliably delivers multiple distinct payloads, enables combinational mixing, and enables point-of-care reconstitution of solid-state payloads.

**Teaser:** Leveraging biomimetic design and advanced 3D printing, we developed high-resolution microfluidic microneedle array patches (MAPs) that overcome payload and scalability challenges, offering a versatile and efficient platform for precise transdermal drug delivery.

## Introduction

Among many technologies, transdermal drug delivery enables safe, painless, and self-administrable drug and vaccine delivery to patients’ skin (*1–4*). Microneedle array patches (MAPs)—including solid, coated, dissolving, and hollow types—are a promising transdermal delivery technology to replace traditional hypodermic injections (*5*). MAPs have been shown to deliver a precise and controlled liquid therapeutic dose and effective transdermal treatment (*6–11*). However, the universal adoption of a MAP technology has been limited. This is because MAP technology has faced challenges including large cargo loading and delivery volumes, treatment formulation versatility, and manufacturing scalability (*12–15*). A universal MAP technology that can reliably and repeatably deliver various payload formulations, from liquid therapeutics to solid-state treatments, at tunable volumes presents a significant unmet need.

Hollow MAPs, which use microfluidic channels to deliver liquid formulations directly into the skin, are effective for rapidly administering large volumes of liquid therapeutics. However, their design constraints and complex manufacturing processes limit their ability to deliver solid-state payloads or multiple therapeutics, which impedes their commercial potential. Integrating microfluidic channels into hollow MAP designs can enhance their functionality and versatility, transforming their use in transdermal delivery and monitoring (*16–22*). Microfluidic technology manipulates small fluid volumes for various operations, including rapid single-payload transport and multi-payload mixing. For instance, microfluidics can simultaneously deliver multiple payloads, reducing treatment time and cost, while facilitating the co-administration of independent biologics, separately, without co-formulation that might interfere with their therapeutic efficacy (*23–25*). Furthermore, microfluidic reservoirs and reconstitution chambers enable MAPs to deliver reconstituted solid, shelf-stable, payloads in a single dose administration, reducing reliance on cold-chain storage and external formulation development (*26, 27*).

Traditional methods for fabricating hollow MAPs, such as micro-molding and photolithography, are labor-intensive and are limited to two-dimensional designs which hinders the scalable integration of microfluidic features with microneedle technology (*1, 4, 28, 29*). The emergence of additive manufacturing (AM) offers a promising alternative. AM allows for the digital design and fabrication of intricate three-dimensional (3D) microfluidic structures alongside microneedles in a single step, avoiding the added costs of complex design, resulting in microfluidic microneedles that are cost-effective and comparable in price, or even less expensive, than traditional microneedle manufacturing methods (*19*). Among AM techniques, digital light processing (DLP) stands out for its scalability and high resolution (*30*). DLP uses two-dimensional XY-projections of ultraviolet (UV) light to cure layers of photopolymerizable resin sequentially. However, resolving negative spaces, such as microfluidic structures, has historically been a challenging due to UV light accumulation in previously printed layers having intended negative spaces, which can lead to overcuring and hinder the fabrication of high-resolution microfluidic MAP devices (*31*).

We recently introduced a new method of vat-based 3D printing that we refer to as Injection Continuous Liquid Interface Production (iCLIP). iCLIP is an advancement of DLP technology that allows for the fabrication of structures with high resolution in all three cartesian coordinates (*33–35*). In this work, we leverage the iCLIP methodology to manufacture high-resolution microfluidic MAPs that were previously unattainable. The iCLIP process relies on mechanically injected resin renewal at the build surface by feeding oxygen-inhibited resin through the build platform and through the microfluidic channels of our device during the printing process (*33, 36*) (**Fig. 1A,B**).

**Fig 1.**
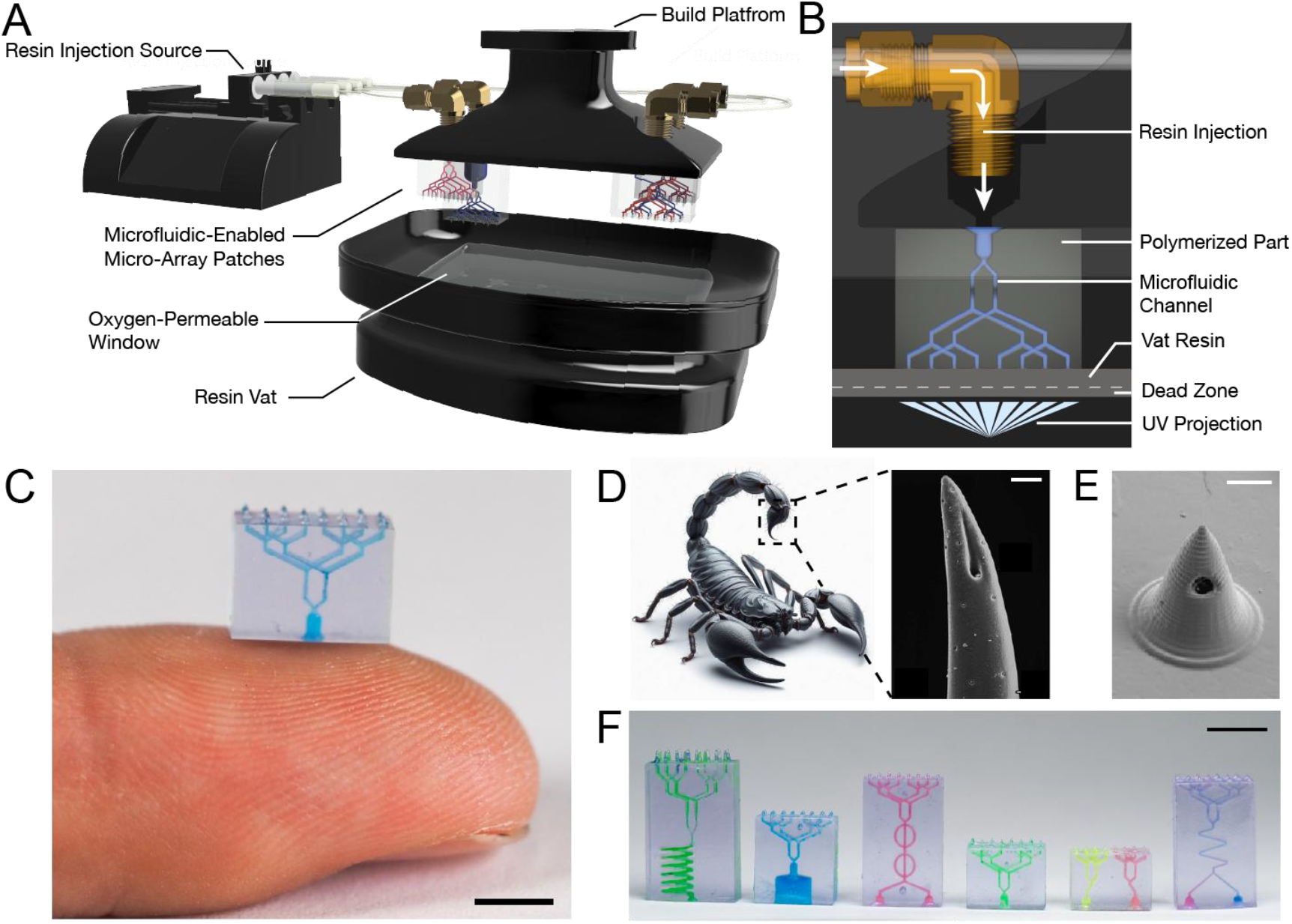
Design of Biomimetic Microfluidic Enabled MAPs. **(A)** Injection CLIP process schematic for microfluidic MAP fabrication. **(B)** Injection CLIP process at the dead zone. **(C)** Photograph of microfluidic MAP on a fingertip. Scale bar 4mm. **(D)** Illustration and scanning electron microscope image of a scorpion stinger (*32*). Scale bar 20 μm. **(E)** 3D printed biomimetic microneedle design. Scale bar 200 μm. **(F)** A diverse array of varying functionality microfluidic enabled MAPs. Scale bar 4 mm.

In this work, we leverage the iCLIP technology to develop a microfluidic MAP technology which enhances the functionality and performance of traditional MAP designs. First, inspired by the design of stingers of insects and the canines of poisonous reptiles, we introduce a biomimetic microneedle design aimed at safeguarding microfluidic inlets from clogging—by strategically positioning the orifice of the needle in a manner that mechanically shields it from clogging during needle insertion into the skin increasing the efficiency of therapeutic MAP delivery (**Fig. 1D, E**). In parallel, we have designed, and 3D printed free-form, non-moldable 3D microfluidic structures, to unite with biomimetic microneedles to rapidly deliver various model therapeutic payloads—including mRNA, proteins, and small molecules—to the transdermal space in a reliable and repeatable manner. As a result, microfluidic MAPs enable one to go beyond simple delivery pathways to co-deliver multiple distinct payloads, perform concurrent mixing and delivery, and reconstitute solid-state products for delivery at point-of-care (**Fig. 1F**).

## Results

### Biomimetic MAP Design Fabrication and Characterization

Venomous animals produce and store venom within a bulb-like reservoir and can rapidly inject it into tissue through an exocuticle stinger. The stinger’s structure has evolved to feature a solid tip with a micropore on the side for venom delivery. Studies have shown that the stinger’s solid tip and side micropore enable efficient, clog-free delivery by preventing obstruction and ensuring consistent injection (*37–40*). In contrast, traditional hypodermic microneedles have a beveled tip with a micropore at the apex. The open tip of a hypodermic needle can compress and fracture when puncturing the stratum corneum or become clogged by dense dermal and epidermal tissue, leading to inconsistent delivery of therapeutic agents (*41–43*). Inspired by the stinger of insects, such as the scorpion, and prior work we have developed a microneedle design that ensures reliable insertion and consistent delivery of payloads (*44*). This design features a solid polymeric needle tip to puncture the stratum corneum (SC) and a 100 μm diameter microfluidic channel within the needle to deliver therapeutics through the side, protecting the microfluidic channel from clogging (**Fig. S1**).

AM processes allow the precise control of microneedle height, enabling targeted delivery to different regions of the transdermal space. In this study, we have designed and fabricated biomimetic microneedles with heights ranging from 600 μm to 900 μm (**Fig. 2A**). All needles were designed to maintain a consistent base diameter of 500 μm, varying only in total height, thus altering their aspect ratio. Each needle height was incorporated into a 16-needle rectangular patch (**Fig. S1**). The iCLIP-based process was then used to print high-resolution microfluidic MAPs with KeySplint Hard resin, often used to make FDA-cleared, biologically compatible dental products (*11*). To assess the ability of the biomimetic MAP to penetrate skin, each MAP was inserted into excised porcine skin using a spring-loaded applicator, followed by applying thumb pressure for 5 seconds (**Fig. S2**). The penetration depth of each microneedle was then evaluated through optical coherence tomography (OCT) and histology. **Figures 2B and 2C** show the SC puncture profile of the 900 μm microneedles, demonstrating uniform penetration into the transdermal space. This procedure was repeated for the other needle heights, with each successfully penetrating the porcine skin at different depths, highlighting the tunable depth-targeting capability of each MAP (**Fig. 2F**)

**Fig 2.**
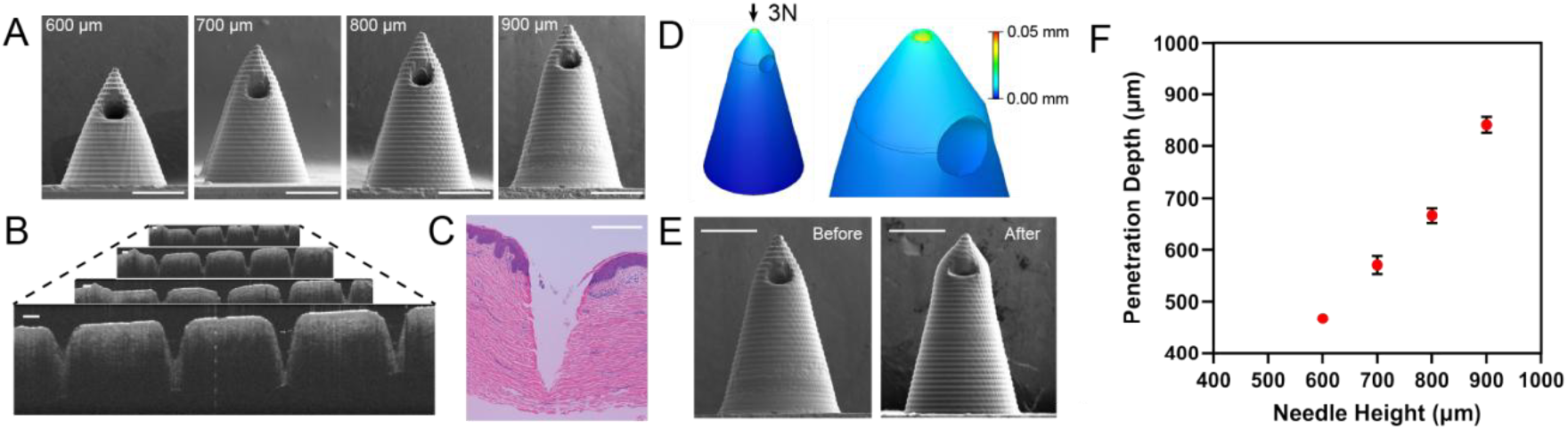
Biomimetic Microneedle Design and Performance. **(A)** Scanning electron microscopy images of biomimetic microneedles with varying heights. Scale bars represent 250 μm. **(B)** Tiled optical coherence tomography (OCT) images showing a 900 μm microneedle array patch (MAP) inserted into excised porcine skin. Four individual OCT scans were stitched together to represent the entire insertion area. Scale bars represent 250 μm. **(C)** Histological cross-section of a 900 μm biomimetic microneedle within excised porcine skin. Scale bars represent 250 μm. **(D)** Finite element analysis of biomimetic microneedle mechanics. **(E)** Scanning electron microscopy images showing biomimetic microneedles before and after insertion. Scale bars represent 250 μm. **(F)** Characterization of penetration depth as a function of microneedle height. All data are presented as mean ± SEM of samples of individual microneedles on MAP.

Next, we examined the mechanical integrity of the biomimetic microneedles to ensure they would not fracture upon insertion into the skin. First, we developed a finite element analysis (FEA) model to simulate microneedle tip deflection after skin insertion. In this model, a 3 N structural force load was applied to each 900 μm microneedle, simulating the force required to puncture the skin (*45*). **Figure 2D** shows the FEA results, predicting minimal tip deflection and no fracture upon insertion. To validate the model, **Figure 2E** compares the needle design before and after the application of a 3 N point load. Additionally, scanning electron microscopy (SEM) images confirm that the 900 μm biomimetic microneedles do not fracture under this force and retain 85 percent of their original tip sharpness. This demonstrates that these microneedles are resistant to failure in compressive environments during skin penetration.

### *In vivo* transdermal delivery of liquid payloads with microfluidic MAPs

We designed a 16- needle microfluidic MAP, which includes a microfluidic network connecting a payload reservoir to 16 independent microneedle outlets, each 600 μm in height, capable of rapidly delivering the payload to the transdermal space (**Figs. 3A, B**). Each microfluidic MAP cartridge is paired with a liquid dispenser that holds and mechanically injects the payload through the microfluidic MAP (**Figs. 3A, B**). After assembly, each microfluidic MAP was loaded with firefly luciferase (FLuc) mRNA ionizable lipid nanoparticles (LNPs) (ALC-0315) and applied to mice using a spring-loaded applicator. **Figures 3C and 3D** demonstrate successful delivery of FLuc mRNA LNPs to mice within 10 seconds, after which the microfluidic MAPs were removed. In this experiment, 23 µL was loaded, and 21 µL was delivered, leaving a residual volume of approximately 2 µL in the device. To measure successful *in vivo* delivery, luciferase protein production from the FLuc mRNA was quantified. D-Luciferin substrate was injected into the mice prior to imaging to detect luciferase protein, with bioluminescence images taken at 5, 24, 48, 72, 96, and 120 hours post-delivery (**Fig. 3C**).

**Fig 3.**
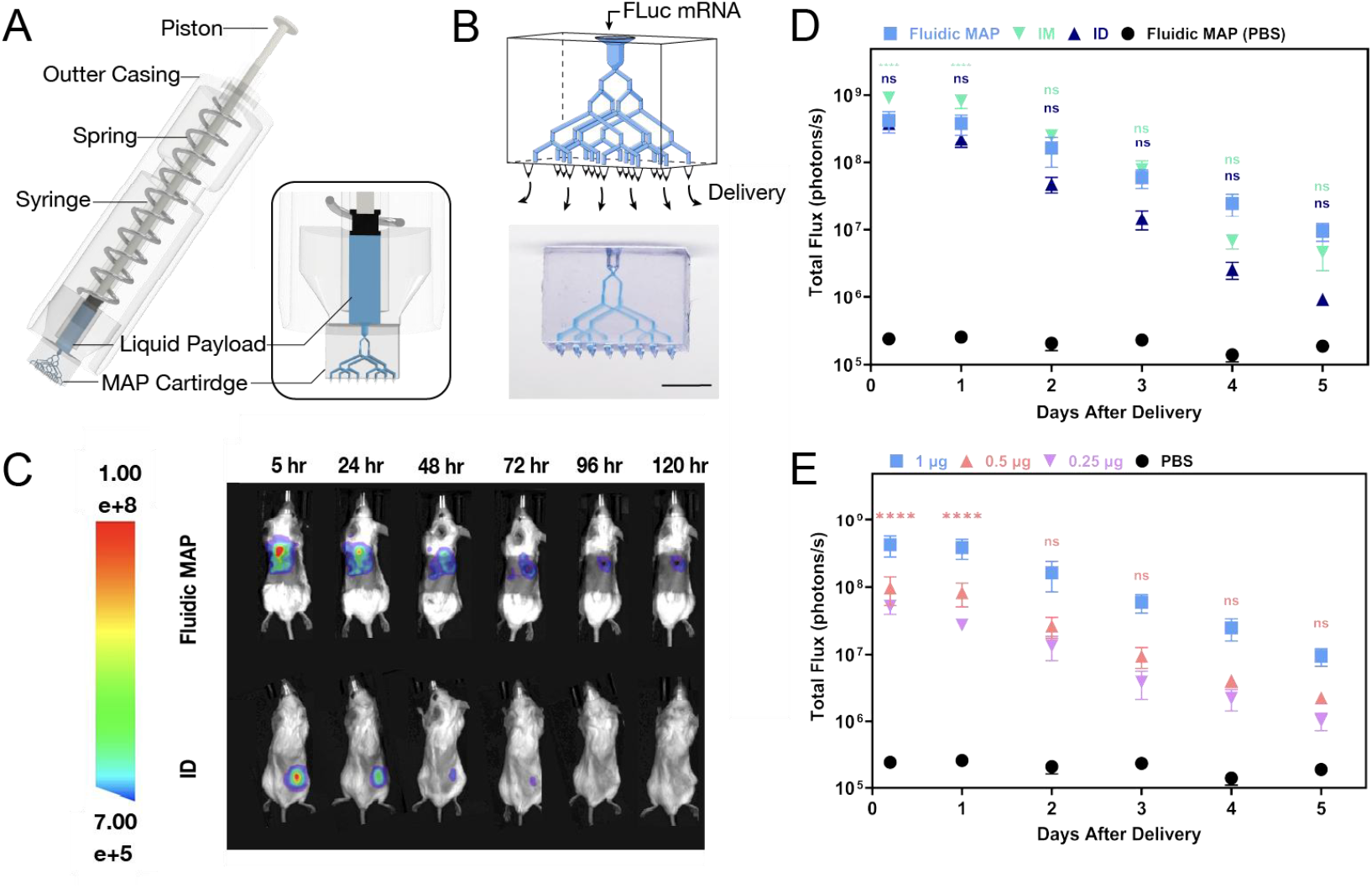
Microfluidic MAP delivery of firefly luciferase (FLuc) mRNA LNPs in mice. **(A)** Schematic of microfluidic MAP assembly, featuring an integrated single-payload applicator. **(B)** Single-payload microfluidic MAP cartridge. Scale bar 4 mm. **(C)** Luminescent images of mice over time following the delivery of 1 µg FLuc mRNA formulated in LNPs with ALC-0315 as the ionizable lipid at the target site. **(D)** Performance quantification of 1 µg FLuc mRNA LNPs in DPBS delivery via microfluidic MAP compared to ID and IM controls (n=5 mice for all groups). **(E)** Dose-dependency quantification for the microfluidic MAP delivery of FLuc mRNA in ALC- 0315 LNPs at doses of 0.25, 0.5, and 1 µg (n=5 mice per condition). All data are presented as mean ± SEM of samples of individual animals. For Fig. D,E, data were analyzed by two-way ANOVA. In Fig. D, the analysis between 1 and 0.5 µg doses is displayed. In Fig. E, the analysis between the microfluidic MAP (1 µg dose) and the IM (1 µg dose) and ID (1 µg dose) are displayed. In both D and E, * P<0.1, ** P<0.01, *** P<0.001, ****P<0.000.

The delivery of liquid FLuc mRNA LNPs using microfluidic MAPs was compared to intradermal (ID) and intramuscular (IM) injections by evaluating the bioluminescent signal from an equivalent dose of FLuc mRNA delivered via these routes. Within 24 hours, microfluidic MAP delivery achieved a comparable level of luciferase expression to both IM and ID injections (**Fig. 3D**). Furthermore, we examined the dose-dependency of the microfluidic MAP technology by loading 0.25, 0.5, or 1 µg of FLuc mRNA in 23 µL, delivering 21 µL with approximately 2 µL of residual volume (**Fig. S3**). As expected, dose-dependent expression profiles were observed (**Fig. 3E**). These results indicate that the microfluidic MAP system can rapidly and precisely deliver a liquid payload at varying concentrations.

### Versatile delivery of liquid payloads with microfluidic MAPs

The ability to co-deliver multiple therapeutics or vaccines simultaneously, intradermally or subcutaneously, with a single administration would be an advantageous dosage form. Such a capability would streamline the administration process and improve patient compliance allowing clinicians to administer several therapeutics simultaneously, eliminating the need for extensive formulation development to ensure compatibility. An example of a drug that may benefit from co-delivery on a single MAP are glucagon-like peptide-1 (GLP-1) receptor agonists (GLP-1Ras) which are being studied for codelivery with other drugs including sodium-glucose co-transporter-2 (SGLT-2) and insulin for enhanced functionality in diabetes management and weight loss (*46–48*). Additionally, cancer drugs such as doxorubicin are being investigated with co-delivery of additional drugs like curcumin to help improve efficacy and reduce toxicity (*49*). There are also applications of co-delivery of antiviral drugs for disease treatment, for example, codelivery of sofosbuvir and ribavirin for Hepatitis C treatment (*50*). To explore the capability of microfluidic MAPs for multi-payload delivery, we designed a 16-needle MAP cartridge with two independent microfluidic networks, each connecting a source port to eight microneedle outlets (**Fig. 4A**). The dual-payload delivery microfluidic MAP cartridge was assembled into the delivery device and applied to a mouse (**Fig. S4**). Through one network, 1 µg of FLuc mRNA in LNPs (in 23 µL DPBS) was loaded and 22 µL was delivered within 10 seconds, leaving a residual volume of approximately 1 µL (**Fig. S3**). Immediately afterward, 45 µg of Texas Red-labeled OVA protein (in 23 µL DPBS) was loaded into the second network, and 22 µL was delivered in another 10-second application, also leaving about 1 µL of residual volume (**Fig. S3**). After both payloads were delivered, the microfluidic device was removed. Bioluminescence and fluorescence images were taken at 5, 24, 48, 72, 96, and 120 hours post-delivery to quantify the transfection of FLuc mRNA and the delivery of Texas Red-labeled OVA protein (**Fig. 4B**). The effectiveness of dual-payload delivery using microfluidic MAPs (FLuc mRNA in ALC-0315 LNPs and Texas Red-labeled OVA) was compared to intradermal injections by analyzing the bioluminescent and fluorescent signals from side-by-side injections of both payloads (**Figs. 4C, D**).

**Fig 4.**
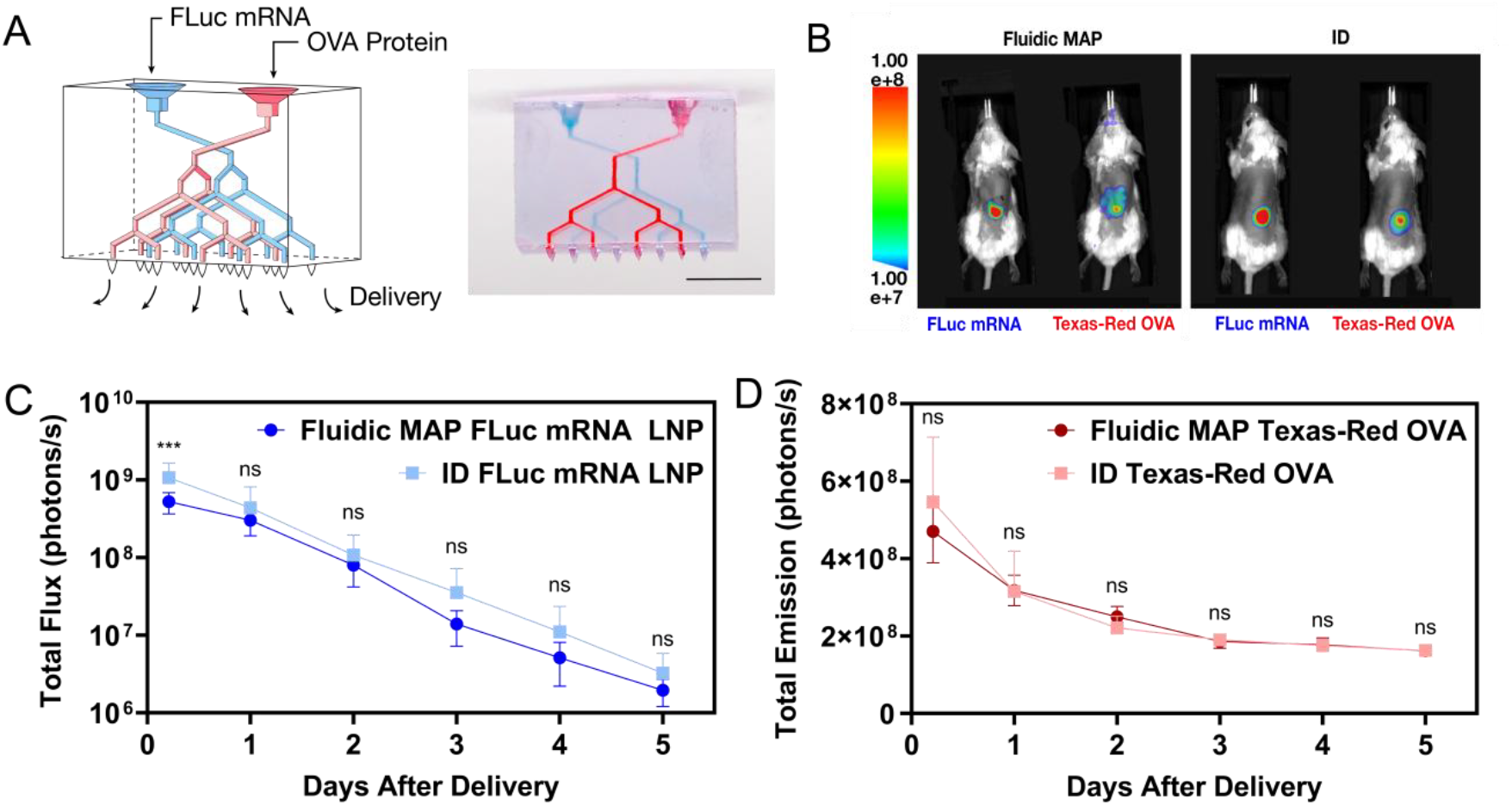
Transdermal delivery of multiple payloads using microfluidic MAPs. **(A)** Dual-payload microfluidic MAP schematic and cartridge. Scale bar 4 mm. **(B)** Luminescent and fluorescent images of mice following the simultaneous delivery of 1 µg FLuc mRNA LNPs in 23 µL and 45 µg of Texas Red-labeled OVA protein in 23 µL in DPBS at the target site with either microfluidic MAP or ID control. Images pictured taken at 5 hours after application. **(C)** Performance quantification of the delivery of 1 µg FLuc mRNA LNPs in 23 µL delivered via microfluidic MAP compared to ID control (n=5 mice per condition). **(D)** Performance quantification of the delivery of 45 µg of Texas Red-labeled OVA protein in 23 µL at Texas Red-labeled OVA protein delivery via microfluidic MAP compared to ID control (n =5 mice per condition). All data are presented as mean ± SEM of samples of individual animals. For C and D, data were analyzed by two-way ANOVA where * P<0.1, ** P<0.01, *** P<0.001, ****P<0.000.

To further demonstrate the liquid delivery capabilities of the microfluidic MAP technology, we designed and manufactured a microfluidic-reactor MAP capable of combining substances at the point of care. This system features a microfluidic chip with two inlets, each supplying a different chemical reactant. These reactants are guided through a microfluidic reactor to form a chemical product, which is then distributed via a microfluidic network across a 16-needle MAP for delivery (**Figs. 5A-C**).

**Fig 5.**
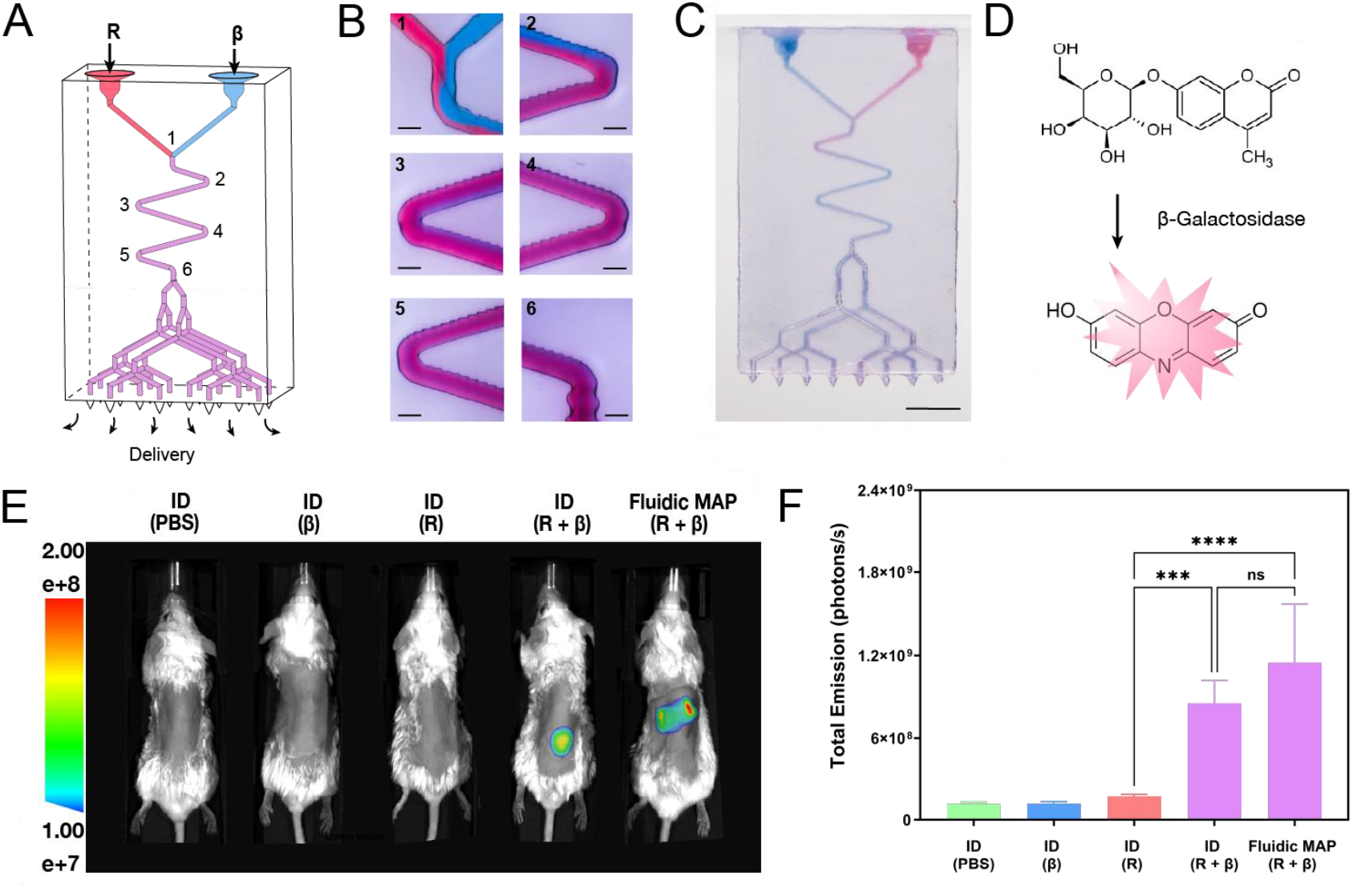
In-vivo transdermal delivery of combined payloads using a mixing microfluidic MAP. **(A)** Mixing-payload microfluidic MAP cartridge schematic. **(B)** Microscopy images of microfluidic MAP mixing unit using two color dye streams. Scale bar 200 μm. **(C)** Mixing-payload microfluidic MAP cartridge. Scale bar 4 mm. **(D)** Reaction of resorufin β-D-galactopyranoside with β-Galactosidase. **(E)** Florescent images of mice over time following the delivery of 6 µg of resorufin β-D-galactopyranoside and 38 µg β-Galactosidase at the target site taken within 15 minutes after application. **(F)** Performance quantification of mixed payload delivered via microfluidic MAP compared to ID controls. For **Fig. f**, data were analyzed by one-way ANOVA. All data are presented as mean ± SEM of samples of individual animals where \**P<0*.*1, ** P<0*.*01, *** P<0*.*001, ****P<0*.000.

To execute this operation, the microfluidic MAP cartridges were assembled into the delivery devices and applied to mice (**Fig. S5**). To demonstrate the chemical reaction capabilities of the mixing cartridge, two reagent payloads — (*1*) resorufin-β-D-galactopyranoside and (*2*) β- Galactosidase — were simultaneously injected into the mixing unit. Inside the unit, an enzymatic reaction occurred, where resorufin-β-D-galactopyranoside was hydrolyzed by soluble β- galactosidase, producing a detectable fluorescent resorufin product (**Fig. 5D**) (*51*). A total of 23 µL of each payload was loaded into the device, resulting in a combined volume of 46 µL, with a residual volume of approximately 8 µL for the product (**Fig. S3**). The fluorescent product was then rapidly delivered through the microfluidic network into the mice. Fluorescence images were captured within 15 minutes of delivery (**Fig. 5E**). As a control, equivalent quantities of resorufin- β-D-galactopyranoside and β-Galactosidase were mixed in an Eppendorf tube by rapid pipetting and injected intradermally into the mice. The fluorescent signals generated from the microfluidic MAP mixing and delivery were compared to those from the intradermal control (**Fig. 5F**).

### Delivery of solid-state payloads with microfluidic MAPs

In addition to liquid payload delivery, we designed a microfluidic MAP capable of containing a solid-state cargo in a reservoir, which can be reconstituted and delivered at the point of care in a single step (**Figs. 6A, B**). This design minimizes the process time and complexity compared with traditional solid-state therapeutic formulations. Specifically, 12 µL (equivalent to 36 µg) of Texas Red-labeled ovalbumin protein in a 20 wt/v percent sucrose solution was lyophilized in the microfluidic reservoir. Studies indicate that ∼80 % of lyophilized protein activity can be retained with optimized formulation conditions (*52–54*). After lyophilization, each cartridge was loaded with 23 µL of PBS as the reconstitution fluid, ensuring the lyophilized cake and PBS separate until application (**Fig. S6**). Upon application, the reconstitution fluid rehydrated the lyophilized cake within 10 seconds, and the rehydrated formulation was then delivered through the microfluidic array. The residual volume in the device was approximately 5 µL. Fluorescence images were taken at 1, 24, 48, 72, 96, and 120 hours following the delivery of the Texas Red-labeled OVA (**Fig. 6C**). To evaluate performance, delivery via the microfluidic MAP was compared to intradermal injections by measuring the fluorescent signals from an equivalent dose of liquid Texas Red-labeled OVA (**Fig. 6D**). Initially, comparable fluorescence signals were observed between the two delivery routes.

**Fig 6.**
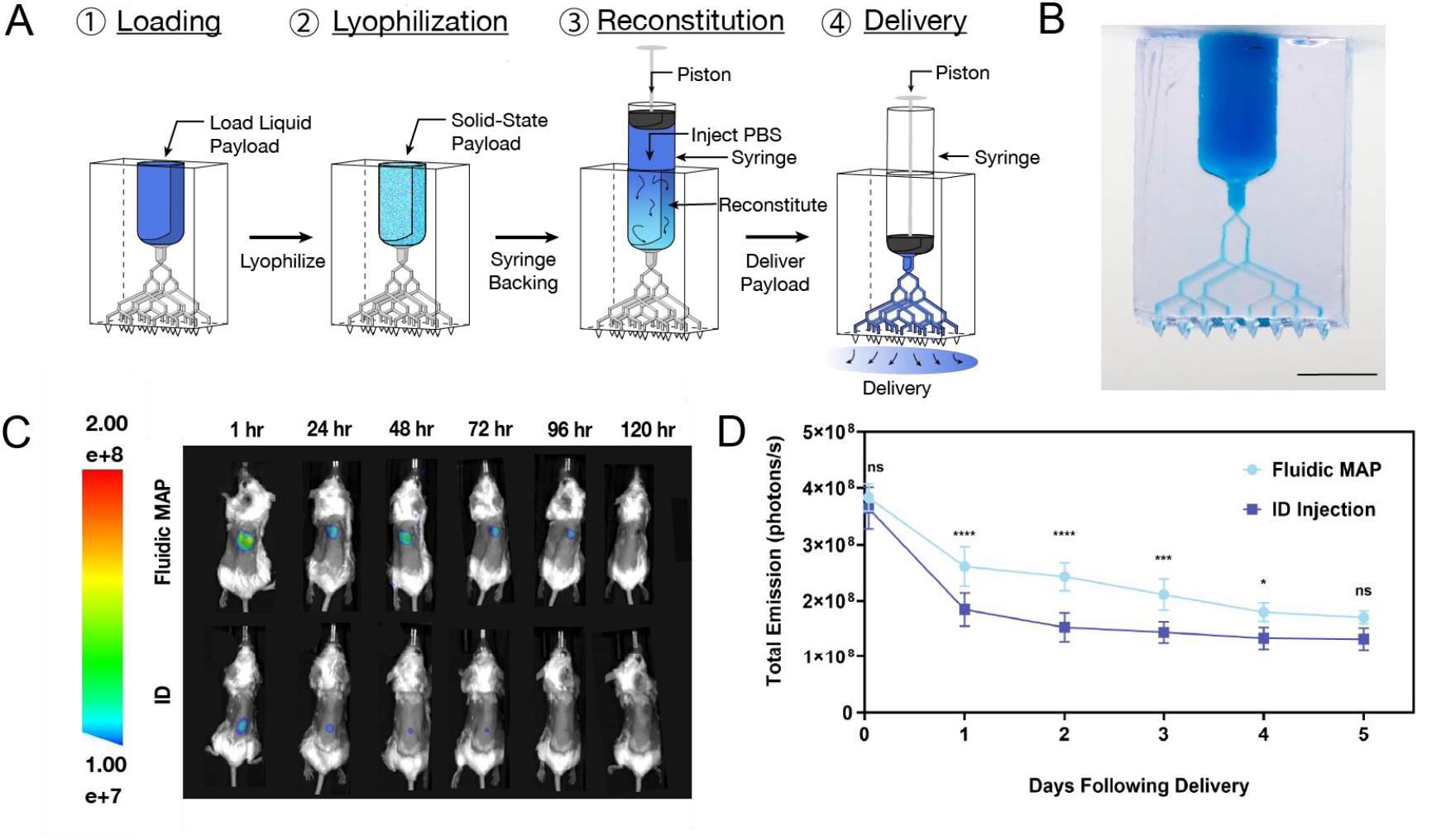
Transdermal delivery of solid-state lyophilized payload using microfluidic MAPs. **(A)** Schematic illustrating the process of lyophilization, reconstitution, and delivery of the solid-state payload. **(B)** Solid-state microfluidic MAP cartridge. Scale bar 4 mm. **(C)** Fluorescent images of mice over time following the delivery of 36 µg of Texas Red-labeled OVA protein at the target site. **(D)** Quantification of the performance of lyophilized 36 µg Texas Red-labeled OVA protein delivery via microfluidic MAP compared to ID control (n=5 mice per group). All data are presented as mean ± SEM of samples of individual animals. For Fig. D, data were analyzed by two-way ANOVA where * P<0.1, ** P<0.01, *** P<0.001, ****P<0.000.

## Discussion

Microneedle array patches (MAPs) have revolutionized drug delivery and diagnostics by enabling minimally invasive, pain-free, and self-administrable treatments. Unlike hypodermic needles, MAPs bypass pain receptors in the dermis, reducing discomfort while enhancing patient compliance. Their small size and ability to deliver drugs transdermally eliminate the risks of needle-stick injuries and biohazardous waste. However, despite these advantages, existing MAP technologies are limited by scalability, complex formulations, and in needle geometry, restricting their widespread clinical adoption (*2, 4, 5*).

Here, we introduce Generation 4.0 MAPs, which overcome these challenges by integrating digitally designed freeform microfluidic networks directly behind the microneedles. This innovation enhances dose precision, payload versatility, and manufacturing scalability, offering an improvement over previous MAP generations.

Generation 1.0 MAPs relied on labor-intensive microelectronic manufacturing, producing simple, silicon-based needles that lacked scalability and increased costs. Generation 2.0 MAPs introduced a broader range of materials and fabrication techniques, such as traditional microfabrication and soft lithography, but still suffered from geometric constraints. Generation 3.0 MAPs overcame these structural limitations by employing high-resolution AM, enabling more complex needle architectures. However, these systems remained limited in functional integration, particularly in fluid handling (*1*).

Generation 4.0 MAPs advance beyond these limitations by integrating a microfluidic delivery platform directly into the MAP structure. This system allows for precise liquid dose delivery within seconds while maintaining the simplicity and efficiency of a patch-based approach. Crucially, these microfluidic MAPs support the simultaneous or sequential administration of two distinct payloads through separate inlets and microfluidic networks—an advancement not achievable with traditional hollow MAPs. This dual-delivery capability offers several critical advantages over current clinical practices. For instance, co-delivery enables the simultaneous administration of multiple vaccines, such as flu and COVID-19, with a single MAP application, significantly improving vaccination efficiency and reducing patient stress (*55, 56*). Additionally, dual-payload administration allows for combination therapies, where an adjuvant can enhance the biological response of a primary therapeutic without requiring extensive formulation modifications. Beyond dual payload delivery, Generation 4.0 MAPs introduce an integrated mixing capability, enabling the controlled combination of two distinct therapeutics just before administration. This eliminates the need for pre-mixed formulations, reducing storage complexity and formulation instability (*57, 58*). The ability to generate unstable products from stable precursors immediately before delivery further broadens the scope of applicable therapies.

One major drawback of hollow MAPs is their inability to deliver dry, thermostable formulations, restricting their use to cold-chain-dependent biologics. In contrast, Generation 4.0 MAPs can integrate lyophilized or air-dried payloads directly within the microfluidic reservoir, enabling temperature-stable biologic storage and reconstitution upon application. This eliminates the need for polymer excipients often required in traditional MAP formulations, streamlining biologic development and preserving therapeutic integrity (*15*). By leveraging established lyophilization protocols, microfluidic MAPs facilitate the integration of temperature-sensitive biologics while avoiding the complexities of cold-chain logistics. Furthermore, the automated reconstitution process minimizes handling errors and reduces contamination risks, a leading cause of sterility loss during self-administration (*59, 60*). While MAPs in this study were used immediately after lyophilization, long-term storage would require a dry, moisture-resistant environment to maintain protein stability (*61*).

In the present study, we investigated: i) liquid payload delivery, ii) dual payload delivery, iii) mixed payload delivery, and iv) lyophilized solid-state payload delivery, using the microfluidic MAP technology *in vivo* with mice and conducted mechanical characterization using *ex vivo* porcine skin. Further research is needed to explore the technology’s applications for specific uses, such as vaccine and therapeutic delivery. Future work will include *in vivo* immunization studies and assessments of the technology’s efficacy in larger animal models and human participants to facilitate translation to clinical practice. Given its potential for standardization and precision across manufacturing, payload loading, and delivery, along with its capability for microfluidic network modification, we anticipate that this technology will become a reliable and reproducible MAP delivery system. Moreover, this platform’s versability shows promise for advancing current clinical therapeutic delivery methods beyond the capabilities of existing MAP technologies.

## Materials and Methods

### Biological Models and Approvals

Twelve-week-old female BALB/c mice (Charles River Laboratories) were utilized for all *in vivo* experiments. All *in vivo* procedures involving animals were approved by the Institutional Animal Care and Use Committee (IACUC) and the Administrative Panel on Laboratory Animal Care (APLAC) at Stanford University before the start of the study (APLAC #34358). For mechanical characterization, excised porcine ear skin was sourced from recently euthanized Yorkshire pigs from the Stanford (APLAC #34192).

### Microfluidic-Enabled MAP Design

Autodesk Fusion (Autodesk) was used to design all microfluidic-enabled MAPs. Each MAP used in this study featured a uniform 16-microneedle configuration, with variations only in microneedle height. The MAPs consisted of a 4 by 4 array of microneedles, spaced 2 mm apart horizontally and 1 mm apart vertically, with every other row indented to maximize spacing between the microneedles (**Fig. S1**). Each microneedle has a 500- μm diameter base, tapering to a 200-μm diameter offset plane, located 200 μm below the microneedle tip. This plane is lofted to a point, ensuring consistent tip geometry across microneedles of different heights (**Fig. S1**). Additionally, each microneedle incorporates a 100- μm microfluidic channel that connects the microfluidic network to a microneedle outlet positioned 200 μm below the tip (**Fig. S1**).

### iCLIP Production of Microfluidic-Enabled MAPs

All microfluidic-enabled MAPs were printed on a prototype high-resolution CLIP printer, the S2, produced by Carbon3D (Carbon3D). The S2 features a pixel size of 5.4 μm and a build area of 14.5 mm by 8.2 mm. To accommodate the iCLIP process, the build platform was adapted to include three microfluidic injection ports capable of supplying fresh resin through the microfluidic networks during printing (**Fig. S7**). Each port is connected to an external Harvard PHD Ultra syringe pump (Harvard Apparatus) to supply fresh resin at a set rate. MAPs were printed with biocompatible KeySplint Hard resin (Keystone Industries).

The printing process was controlled as follows. To ensure strong bonding between the build platform and the 3D part, the initial exposure dose was set to 10 times the minimum required exposure dose to polymerize the first slice layer. After the initial exposure layer, fresh resin was injected through the build platform into the developing negative space at a set flow rate. Specifically, the flow rates for each MAP type were as follows: single payload MAP – 3.0 μL/min, dual payload MAP – 1.0 μL/min, microreactor MAP – 1.0 μL/min, and reservoir MAP – 5.5 μL/min. Subsequent print layers were exposed to 1.5 times the minimum required dose to polymerize each 25-μm layer. After printing was complete, resin injection was stopped, and the 3D parts were cleaned. Isopropyl alcohol was used to clean each printed sample, ensuring that all uncured resin was removed. Finally, the objects were cured in an APM LED UV Cube (APM Technica) for 10 minutes.

### Examining Penetration Depth of Biomimetic Microneedle

Excised porcine skin from the outer ear of euthanized Yorkshire pigs was prepared by removing all hair. The skin was tensioned on a disposable dissection board (Avantor). Each MAP was placed on the surface of porcine skin, and a custom applicator was used to apply approximately 20 N of force for proper contact. Additional thumb pressure was applied to each MAP to ensure full insertion into the skin for 5 seconds. Once applied, MAPs were secured in place with Tegaderm (3M). The skin was then fixed overnight in 10 percent formalin (Millipore Sigma). The following day, the MAPs are removed and imaged using optical coherence tomography (OCT) (Thorlabs). Two-dimensional (2D) slices were analyzed to assess skin penetration depths at the individual microneedle levels (N=3 patches, 16 microneedles per patch, varying microneedle heights from 600 μm to 900 μm) for each row of skin penetration sites. These penetration depths were averaged, and the standard error was reported as the error bars in **Figure 2E**.

### Examining Mechanical Strength of Biomimetic Microneedle

Finite-element-analysis (FEA) of the 900-μm tall biomimetic microneedle design was conducted using Fusion 360 (Autodesk), The analysis was specified to reflect physical properties of KeySpint Hard resin, with the following parameters: Young’s Modulus of 2.35 GPa, Poisson’s Ratio of 0.30, shear modulus of 1600 MPa, and density of 1.1 g/cm^3^. A 3 N point load was applied at the tip of each microneedle, and the resulting displacement (mm) of each element was measured to ascertain tip deflection and ultimately microneedle failure.

To simulate skin puncture, a 3 N load was applied to each microneedle on a 16-microneedle array, using an Instron equipped with compression plates (Instron). This load was precisely applied to the tip of each microneedle. The microneedles were then imaged using an Apreo S LoVac scanning electron microscope (SEM) (ThermoFisher) both before and after the load was applied to visualize any deflection or failure.

### Microfluidic-Enabled MAP Delivery Device

Each microfluidic-enabled MAP delivery device consists of four key components: the MAP cartridge, the MAP collar, the payload holder with piston, and a spring-loaded applicator. The MAP cartridge and collar were fabricated on the Carbon M2 printer. The MAP collar functions to connect the syringe to the MAP cartridge, while the spring-loaded applicator applies constant and reproducible force application across MAP replicates (**Fig. S2**).

To assemble each device the microfluidic-enabled MAP is attached to the MAP collar using ultra- violet adhesive (Loctite IND 405). Next, the desired volume of payload is loaded into a syringe and all air is removed. This syringe is then connected to the MAP/collar unit. This delivery device is placed into the spring-loaded applicator and applied to the mice. For all mouse studies, the mice were positioned on their sides, and their skin was stretched before application of the MAPs (*62*) (**Fig. S2**).

### Formulation of FLuc mRNA-LNPs and Liquid Delivery with Microfluidic-Enabled MAP

FLuc mRNA (L-7202, TriLink) was encapsulated in ALC-0315 LNPs using the LipidLaunch™ LNP-0315 Exploration Kit (35426, Cayman Chemicals). The lipids in the kit (ALC-0315, 1,2- DSPC, cholesterol, and ALC-0159) were each dissolved in 200-proof ethanol (Fisher BioReagents) as specified in the kit to make stock solutions. Immediately before complexing with FLuc mRNA, the lipids were mixed into a combined ethanol phase according to the ratios provided in the kit. FLuc mRNA was dissolved in the proprietary Precision Nanosystems Inc. formulation buffer at a concentration of 0.105 μg/μL prior to complexing with the lipids. The FLuc mRNA-LNPs were formulated using a microfluidic mixer (Ignite, Precision Nanosystems Inc.) at a ratio of 3:1 aqueous:ethanol. After mixing, the formulated FLuc mRNA-LNPs were diluted in Dulbecco’s phosphate buffered saline (DPBS) (Corning, Sigma Aldrich). The total volume (50 mL) was divided equally into four centrifugal filter tubes (UFC9100, Amicon 100kDa), and spun at 2000 x g for 30 minutes at 4 °C. Following centrifugation, the concentrated FLuc mRNA-LNPs were recovered from the filter tubes, and sucrose solution was added to bring the final sucrose concentration to 10 wt/v percent. The quantity of mRNA and the encapsulation efficiency of the LNPs were quantified using the QuantiFluor Assay (Promega). Two sets of standards and samples were prepared in either Tris-EDTA buffer obtained from Promega to quantify encapsulated RNA or 2 v/v percent Triton X-100 (Sigma Aldrich) to quantify total mRNA. Size (nm) and polydispersity index (PDI) were measured using dynamic light scattering (DLS) (Zetasizer, Malvern Panalytical). The FLuc mRNA-LNPs were then stored at −80 °C. The FLuc mRNA LNPs were diluted to a concentration of 50 ng/µL in DPBS, and 23 µL were loaded into the MAP application devices for each replicate. Individual devices and microfluidic-enabled MAPs were assembled for each mouse.

Mice for the studies were prepared by shaving the hair from the back and applying Nair depilatory cream (Church and Dwight) for three minutes for complete fur removal the day before the microfluidic MAP applications. Prior to microfluidic MAP application, the microfluidic MAPs were sterilized under UV for 10 minutes. Mice were anesthetized using isoflurane (5 percent induction and 3 percent for maintenance) and their skin was sterilized using an alcohol swab.

To determine dosing for intradermal and intramuscular controls, the residual volume in a microfluidic-enabled MAP device was quantified (as described below), and controls were dosed with the total volume delivered by a microfluidic-enabled MAP minus the residual volume. For example, if 23 µL was delivered with a microfluidic-enabled MAP and the residual volume was 1.5 µL, then 21.5 µL would be delivered for the controls.

To quantify the delivery and expression of FLuc mRNA in mice via microfluidic-enabled MAPs, or intradermal (ID) or intramuscular (IM) controls, bioluminescence was measured using the SII Lago-X IVIS (*in vivo* imaging system) (Spectral Instruments) with medium binning and a 10- second exposure time. Mice were injected intraperitoneally (IP) with 100 μL of D-Luciferin (Research Products International) at a concentration of 50 mg/mL in DPBS and imaged 20 minutes after injection. Bioluminescent signals were quantified using the Aura software (Spectral Instruments). Mice were imaged at 5, 24, 48, 72, 96, and 120 hours following delivery to follow FLuc expression over time for each delivery route.

### Texas Red-Labeled OVA and FLuc mRNA-LNP Preparation for Dual Payload Delivery with Microfluidic-Enabled MAP

Solutions for dual-payload delivery using microfluidic-enabled MAPs were prepared for both Texas Red-labeled ovalbumin and FLuc mRNA LNPs. Texas Red-labeled OVA was dissolved in DPBS at a concentration of 5 μg /μL, and then diluted in DPBS to obtain a delivery mass of 30 μg of Texas Red-labeled OVA in 12 μL of solution. The solution was loaded into one of the two inlets of the device. FLuc mRNA-LNPs were prepared as previously described and diluted to 100 ng/µL in DPBS. A volume of 12 μL of the FLuc mRNA-LNP solution was loaded into the other inlet of the microfluidic-enabled MAP. For each mouse, a delivery device containing both payloads were prepared.

The delivery and expression of FLuc mRNA were quantified using the SII Lago-X IVIS, following the same protocol used for single-payload liquid delivery with microfluidic-enabled MAP. For Texas Red-labeled OVA, quantification was performed using the same protocol as for lyophilized Texas Red-labeled OVA with the SII Lago-X IVIS. Prior to imaging, the delivery site on the mouse’s back was cleaned with an ethanol wipe. Mice were imaged at 5, 24, 48, 72, 96, and 120 hours following delivery of lyophilized Texas Red-labeled OVA.

### Resorufin β-D-galactopyranoside and β-Galactosidase Preparation for Mixing Payload Delivery

Solutions of resorufin β-D-galactopyranoside (Sigma-Aldrich) and β-galactosidase (Sigma-Aldrich) were prepared to demonstrate the mixing of two payloads using microfluidic-enabled MAPs. Resorufin β-D-galactopyranoside was prepared at a concentration of 0.3 mg/mL in phosphate-buffered saline (PBS 1X). β-Galactosidase was dissolved in PBS at a concentration of 1000 units/mL (2 mg/mL). A volume of 23 μL of the resorufin β-D-galactopyranoside solution was loaded into one of the two inlets of the device, while an equal volume of the β-galactosidase solution was loaded into the other inlet. Individual devices with both payloads and microfluidic-enabled MAPs were assembled for each mouse. The two solutions were simultaneously injected into the mixing units, allowing them to react and form the fluorescent resorufin, which was then delivered into the mice.

To quantify the delivery of resorufin to mice via microfluidic-enabled MAPs or ID controls, fluorescence signals were measured using the SII Lago-X IVIS with medium binning and a 10- seconds exposure time. The excitation was set to 570 nm and the emission was set to 610 nm. Fluorescent signals were quantified using Aura software. After delivery, the site on the mouse’s back was cleaned with an ethanol wipe before imaging.

### Texas Red-Labeled OVA Preparation, Loading, and Lyophilization and Delivery with Microfluidic-Enabled MAP

Texas Red-labeled ovalbumin (OVA) (O2302, Thermo Fisher Scientific) was dissolved in DPBS to a concentration of 5 mg/mL. A stock solution of 90 wt/v percent sucrose in DPBS was prepared and an appropriate volume was mixed with the Texas Red-labeled OVA solution to achieve a final sucrose concentration of 20 wt/v percent. A 12 μL aliquot of the solution was loaded into the reservoirs of 5 microfluidic-enabled MAPs. The filled MAPs were placed in a tray lyophilizer (Labconco) and frozen at −55 °C for 3 hours before being dried for 20 hours. After drying, the MAPs were removed from the lyophilizer, assembled into the MAP delivery device, and applied in vivo to mice. To reconstitute the lyophilized payload, 23 μL of DPBS was loaded to each device, which was then used to deliver the reconstituted lyophilized cake to mice (**Fig. S6**). Individual devices and microfluidic-enabled MAPs were assembled for each mouse. Following delivery of lyophilized Texas Red-labeled OVA to the mice, the skin where the MAP was applied was wiped with an ethanol wipe to remove any Texas Red-labeled OVA from the skin surface.

To quantify the delivery of lyophilized Texas Red-labeled OVA to mice via microfluidic-enabled MAPs or ID controls, the fluorescence signal was measured using the SII Lago-X IVIS with medium binning and a 10-second exposure time. The excitation was set to 570 nm and the emission was set to 610 nm. Fluorescence quantification was performed using the Aura software. Prior to imaging, the delivery site on the mouse’s back was cleaned with an ethanol wipe to remove any surface residue. Mice were imaged at 1, 24, 48, 72, 96, and 120 hr following delivery of lyophilized Texas Red-labeled OVA.

### Quantification of Residual Volume in Microfluidic-enabled MAPs

The residual volume (dead volume) in each microfluidic-enabled MAP was quantified using both FLuc mRNA-LNPs and methylene blue dye (Sigma Aldrich). For single-payload liquid delivery, microfluidic-enabled MAPs were applied to mice to deliver FLuc mRNA-LNPs. After delivery, the remaining fluid in the MAP was flushed with 100 μL of DPBS. The QuantiFLuor assay with 2 v/v percent Triton X- 100 was then used to quantify the remaining mRNA in the MAPs. By comparing the concentration of delivered mRNA, the volume of residual fluid in the microfluidic-enabled MAPs was calculated (**Fig. S3**).

Residual volume for single-payload liquid delivery was also calculated by using methylene blue dye. A stock solution of methylene blue dye was prepared in DPBS at a concentration of 5 μg /μL, and 23 μL of this dye was delivered through a microfluidic-enabled MAP. The residual dye in the microfluidic-enabled MAP was then flushed out using 100 μL of DPBS. The concentration of methylene blue dye in the flushed volume was quantified through absorbance measurements. A standard concentration curve was generated for methylene blue, absorbance measured at 660 nm. By interpolating the absorbance measurements of the flushed volumes against the standard curve, the concentration of methylene blue was determined. The residual volume was calculated by comparing the measured dye concentration with the known concentration of the original solution. Absorbance measurements were collected using a plate reader (Spark, Tecan Life Sciences) (**Fig. S3**).

For the solid-state delivery device, 12 μL of methylene dye in 20 w/v percent sucrose was added to the reservoir and lyophilized. To reconstitute the dye for delivery, 23 μL of DPBS was used. The residual dye in the microfluidic-enabled MAPs was flushed out with 100 μL of DPBS and quantified following the previously described protocol (**Fig. S3**).

For the dual-payload delivery device, 23 μL of methylene blue dye was first delivered through one inlet, followed by flushing the residual dye with 100 μL of DPBS. This process was then repeated for the second inlet. Delivering dye through each inlet and subsequently flushing both inlets allowed for quantification of the residual volume in each set of channels within the dual-payload device. The concentration of methylene blue dye in each set of channels was measured using the absorbance method described above, and the residual volume for each set of channels was calculated (**Fig. S3**).

Similarly, for the mixing payload delivery device, 23 μL of methylene blue dye was delivered through one inlet, followed by flushing the residual dye with 100 μL of DPBS. This process was then repeated for the other inlet. The residual volume in each set of channels was determined using the same methylene blue as quantification method described earlier (**Fig. S3**).

## Acknowledgments

This work is supported by the Bill and Melinda Gates Foundation, Grant MAP 3.0 Research and development, INV-046940. IAC and MMD are supported by this grant. Part of this work was performed at the Stanford Nano Shared Facilities (SNSF), supported by the National Science Foundation under award ECCS-2026822. We would also like to acknowledge the Stanford Center for Innovation in In vivo Imaging (SCi3) small animal imaging facility. This work was also supported by the Stanford Soft & Hybrid Materials Facility. We acknowledge the use of DALL·E 3, an AI model, for the development of the scorpion image in Fig. 1e.

## Funding

Bill and Melinda Gates Foundation INV-046940 (IAC, MMD, NUR, GL, DI, YX, MTD, GBJ, JLP, ST, JMD)

## Author contributions

Conceptualization: IAC, MMD, GL, MTD, GBJ, JLP, ST, JMD.

Methodology: IAC, MMD, NUR, GL, DI, YX, MTD, GBJ, JLP, ST, JMD.

Experimental Investigation: IAC, MMD, NUR.

Funding acquisition: MTD, GBL, JLP, ST, JMD.

Project administration: MTD, GBJ, JLP, ST, JMD.

Writing – original draft: IAC, MMD.

Writing – review & editing: IAC, MMD, NUR, GL, DI, YX, MTD, GBJ, JLP, ST, JMD.

## Competing interests

JMD declares that he has an equity stake in Carbon Inc., which is a venture-backed start-up company that owns related U.S. Patent 9,216,546, U.S. Patent 9,360,757, and others. IAC, MMD, NUR, MTD, DI, YX, GBJ, JLP, ST and JMD all have an equity stake in PinPrint, which is an early-stage vaccine and drug delivery company that has licensed MAP technology from Stanford University, the University of North Carolina at Chapel Hill and Carbon. Methods for negative space preservation iCLIP and microfluidic microneedle devices are the subject of a pending patent application. The authors declare that they have no other competing interests.

## Data and materials availability

All data are available in the main text or the supplementary materials.

## Supplementary Materials

Supplementary Figs. S1-S8.

