## Supplemental Information for "Free-Form Microfluidic Microneedle Array Patches"

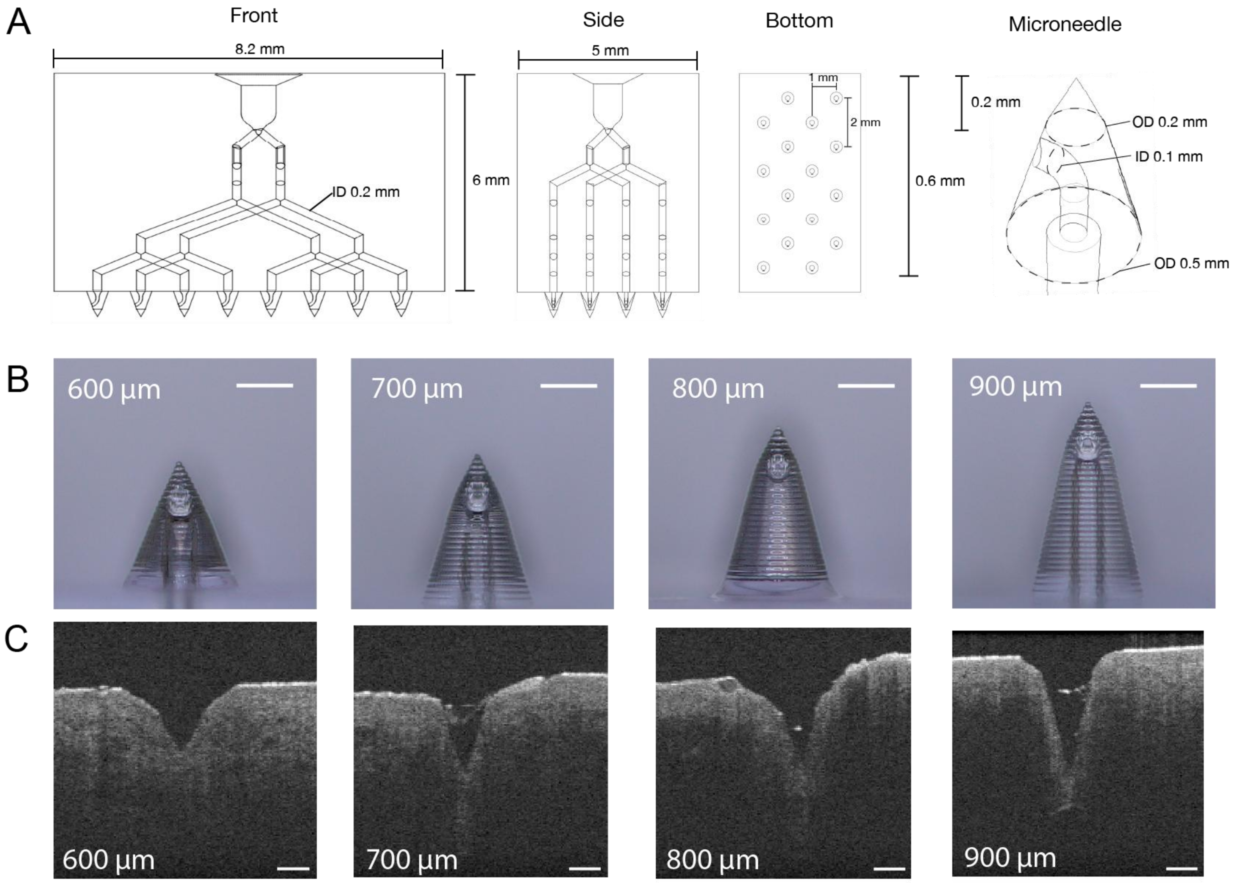


**Fig. S1. Microfluidic MAP cartridge investigation. (A)** Microfluidic MAP dimensioning. **(B)** Brightfield microscope images of microfluidic MAP needles. **(C)** Optical Coherence Tomography images of microfluidic MAP microneedles inserted into excised porcine skin. All scale bars 250 µm.

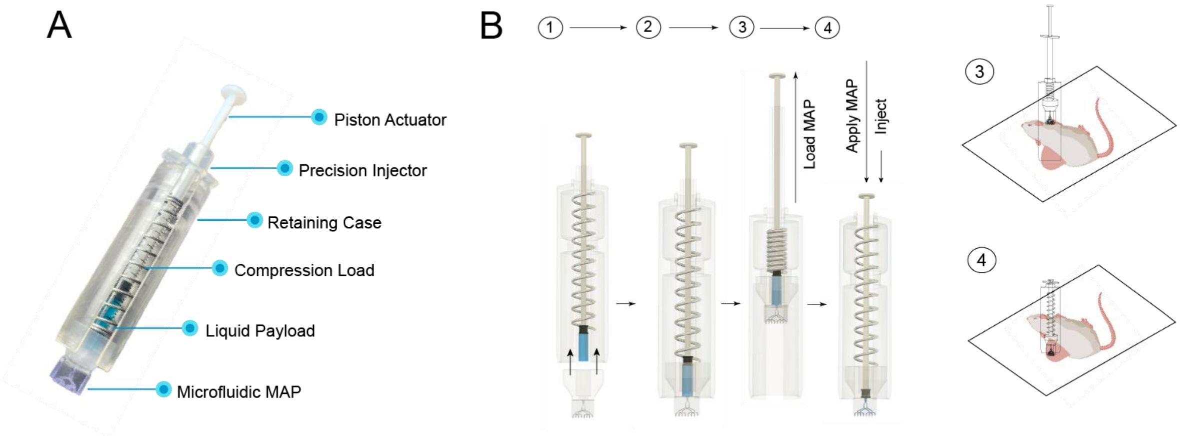


**Fig. S2. Single payload microfluidic MAP device assembly and application. (A)** Single-liquid payload MAP device assembly. **(B)** Single-liquid payload MAP device assembly, application, and delivery schematic.

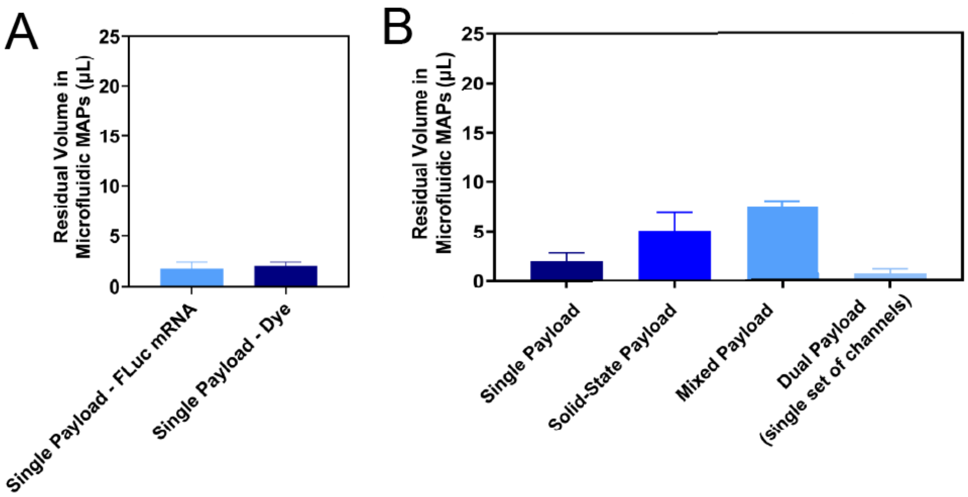


**Fig. S3. Residual Volume in Devices After Delivery. (A)** Residual volume in a single payload device was measured following the delivery of FLuc mRNA and methylene blue dye. **(B)** Residual volume in a single payload, solid-state payload, mixed-payload, and dual-payload (single set of symmetrical channels) was measured using methylene blue dye. For Fig. a, five devices were used for each measurement. For Fig. b**,** five devices were used for the single payload, solid-state payload, and mixed-payload, and 10 devices were used for the dual-payload (single set of symmetrical channels).

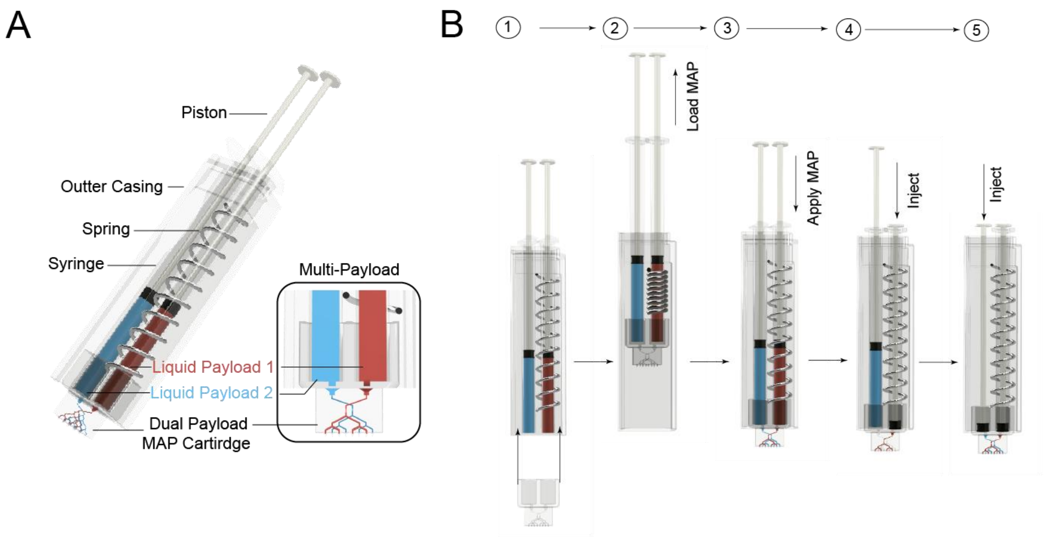


**Fig. S4. Dual payload microfluidic MAP device assembly and application. (A)** Single-liquid payload MAP device assembly. **(B)** Dual-liquid payload MAP device assembly, application, and delivery schematic.

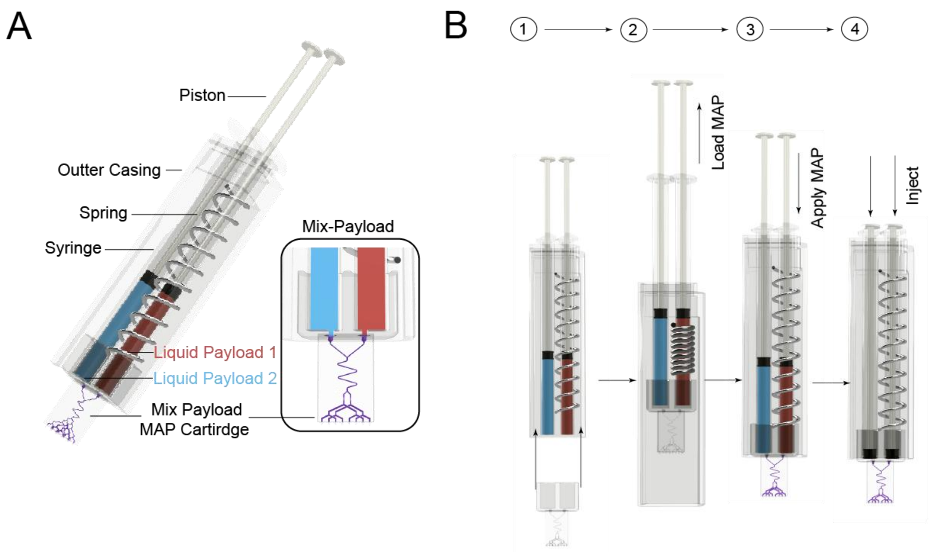


**Fig. S5. Mixing payload microfluidic MAP device assembly and application. (A)** Single-liquid payload MAP device assembly. **(B)** Mixing-liquid payload MAP device assembly, application, and delivery schematic.

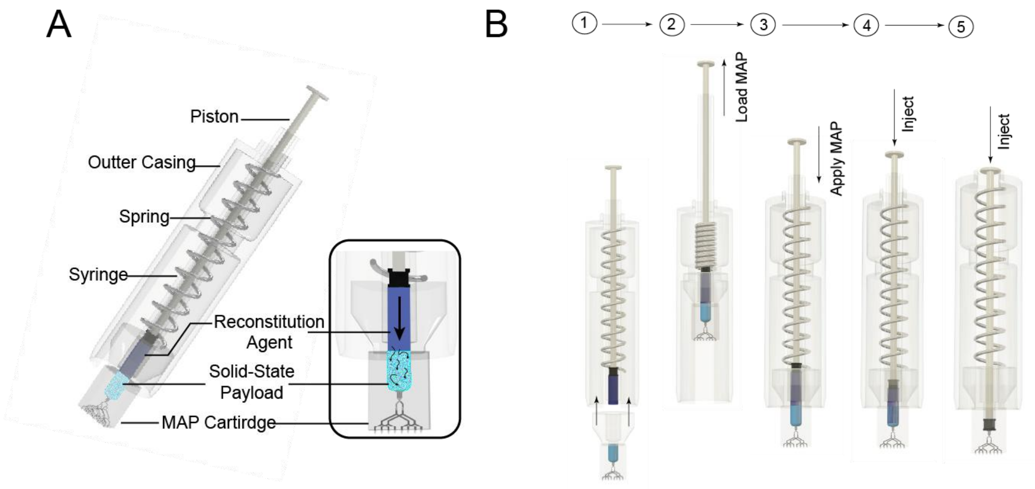


**Fig. S6. Solid-state payload microfluidic MAP device assembly and application. (A)** Single-liquid payload MAP device assembly. **(B)** Solid-state payload MAP device assembly, application, and delivery schematic.

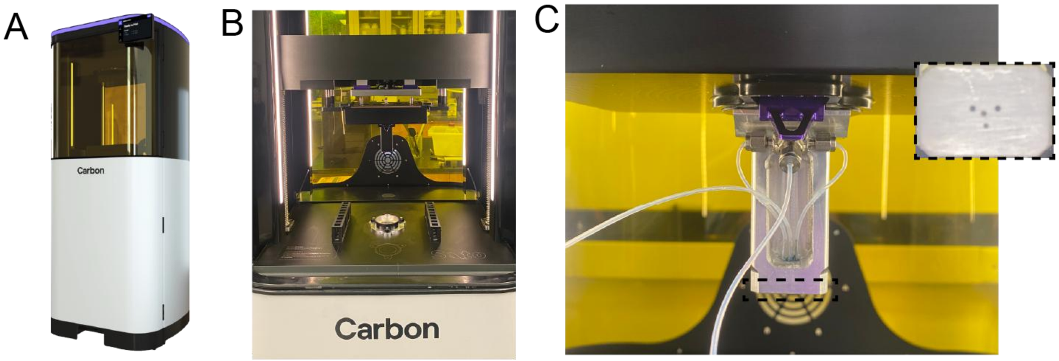


**Fig. S7. Carbon S2 High Resolution iCLIP Printer. (A)** Carbon S2 printer. **(B)** Carbon S2 print area**. (C)** iCLIP build platform with machined injection ports. Inset is a magnified image of this injection ports.

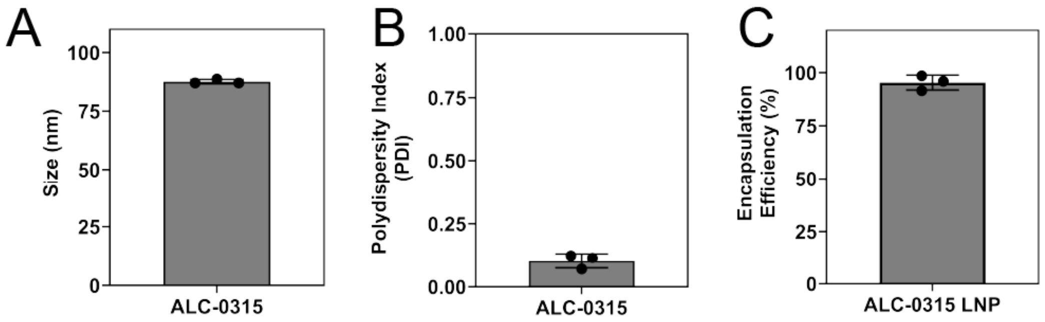


**Fig. S8. FLuc mRNA LNP (ALC-0315) Characterization. (A)** Dynamic light scattering (DLS) measurement showing average hydrodynamic diameter of lipid nanoparticles. **(B)** Polydispersity index (PDI) was measured using DLS. **c,** The encapsulation efficiency percentage of mRNA in the lipid nanoparticles was measured using the QuantiFluor assay. For Fig. a,b, each data point represents one sample, and each data point is an average of at least 10 readings on the DLS, resulting in the average hydrodynamic diameter and polydispersity values displayed on the graphs. For Fig. c, three independent samples were measured. All measurements are shown as mean with SEM.
